# CSTA: A Biomarker Regulating Esophageal Stricture Epithelial Barrier, Potential Medication Target

**DOI:** 10.1101/2023.10.23.563568

**Authors:** Fang Yang, Yiwei Hu, Zewen Shi, Haozhou Jin, Kefeng Hu, Guoliang Ye, Qian Pang, Ruixia Hou, Keqi Tang, Yabin Zhu

**Author notes:** **Correspondence:** Keqi Tang, Institute of Mass Spectrometry, School of Material Science and Chemical Engineering Ningbo University Ningbo 315211, P. R. China, Yabin Zhu Health Science Center Ningbo University Ningbo 315211, P. R. China.

## Abstract

Esophageal strictures resulting from ESD/EMR in the esophagus are distressing, with uncertain causes and treatments due to epithelial barrier dysfunction. We explored cystatin A (CSTA) as a novel biomarker for the esophageal epithelial barrier. CSTA regulates the barrier and is influenced by tacrolimus (FK506). Analyzing biopsy samples by proteomics, we identified CSTA as an esophageal epithelial biomarker. Knocking down CSTA reduced responsiveness to triamcinolone acetonide (TA) and lowered expression of proteins related to cell-cell junctions and barrier function during inflammation. These findings suggest that down-regulate CSTA weakens tight junctions and the esophageal barrier, leading to strictures. Notably, FK506 showed promise in rescuing CSTA-knockdown cells by increasing the expression of proteins related to cell-cell junctions and barrier function. This highlights FK506 as a potential therapy for esophageal strictures in patients with CSTA mutations.

## 1. Introduction

Esophageal strictures resulting from pathological conditions or endoscopic submucosal dissection (ESD)/endoscopic mucosal resection (EMR) surgery due to the malignant tumor are challenging for patients and clinicians. The literature reported that the resection of more than 3/4 circumference of the esophageal mucosa after ESD surgery is a high-risk factor for esophageal stricture, with an incidence of 100% and 56-76% for esophagotomy and non-esophagotomy, respectively(Wang, 2022; Landin & Guerrón, 2020; Shi *et al*, 2014; Kadota *et al*, 2016; Takahashi *et al*, 2015; Yamaguchi *et al*, 2011; Yeung & Chiu, 2017; Maxton *et al*, 1987). Current treatments in the clinic, for example, expansion stents or/and antibacterial drugs, for patients with esophageal strictures are not as effective as they could be since the pathogenesis of the disease is not clear, and the administration is limited.

In recent years, the biological mechanisms leading to esophageal strictures are becoming more perceptible with the development of genomics and proteomics. Some scholars believe that dysfunction of the esophageal epithelial barrier may be one of the initiating causes of esophageal strictures(Blevins *et al*, 2018; Kleuskens *et al*, 2022). After ESD/EMR, the resected mucosal epithelium does not provide a good barrier function, leading to a constant attack of harmful substances on the underlying tissues to cause an excessive inflammatory response and then excessive fibrosis(Iwata *et al*, 2021; Ohki *et al*, 2006). Excessive fibrosis is the most familiar disease mechanism of esophageal stricture(El-Asmar *et al*, 2013; Kataoka *et al*, 2015; Hashimoto *et al*, 2011; Nagami *et al*, 2018). Therefore, studying the epithelial barrier dysfunction in esophageal stricture is essential and critical. Reestablishing the intact esophageal epithelial barrier function as soon as possible can promptly intervene in the occurrence and progression of esophageal stricture. But there are still many mysteries that remain to be solved.

The clinic treatments for esophageal strictures include physical dilatation through esophagoscopy together with local injections or/and oral therapy with glucocorticoid medicines like triamcinolone acetonide (TA) (Nagami *et al*, 2018; Kato *et al*, 2018; Oyama, 2015; Shibagaki *et al*, 2018). However, many patients are insensitive or ineffective to glucocorticoids due to individual differences (Horváth *et al*, 2023; Xia *et al*, 2023; Khlevner *et al*, 2023). Repeated physical expansion is not able to cure the stricture. And it makes patients suffer pain and high economical cost. What the reason causing insensitivity or ineffectiveness to glucocorticoids and whether any other medications could solve the current problem is precisely what we want to explore.

In the human body, the oral mucosa is one of the few tissues capable of perfectly re-epithelialization without scar formation(Roh *et al*, 2017; Griffin *et al*, 2022). Moreover, there are some similarities between oral and esophageal mucosal epithelium. Both are nonkeratinized and complex squamous epithelium. This is a good innovation point when the biomolecules exist in both tissues but differ in protein expression. By comparing the data from proteomics and transcriptomics, we identified a biomolecule, cystatin A (CSTA), expressed in both oral and esophageal tissue. It has never been reported in the literature.

CSTA is a member of the type 1 cysteine protease inhibitors, acid cysteine protease inhibitors, keratolinin, or epidermal SH-protease inhibitors(Duivenvoorden *et al*, 2017). It is also named stefin A and was initially identified as a component of the cornified cell envelope in the keratinocyte. It has been reported to play a role in maintaining the morphology of the cornified envelope as a precursor protein and was proposed as a diagnostic tool for head, neck, and chest cancer and prognosis(Blaydon *et al*, 2011; Moosbrugger-Martinz *et al*, 2015; Wei *et al*, 2022; Gupta *et al*, 2015; Candi *et al*, 2005). Recent researchers show that CSTA plays an essential role in the barrier function of epithelial cells and is mainly expressed in the esophagus, compared with other tissues in the human body. Therefore, CSTA is essential in forming cornfield envelope in keratinocytes, maintaining the epithelial barrier, and promoting epithelial cell development.

In the esophagus, mutations in CSTA may result in the inability of esophageal epithelial cells to take effects properly, which in turn impairs the esophageal epithelial barrier (EEB) function. Incomplete EEB function leads to dysfunction of the esophageal epithelial barrier. Barrier dysfunction allows harmful substances to continuously attack the submucosal tissues of the esophagus, leading to an inflammatory response. Once inflammation cannot be cured in time, the inner cells and tissues will be continuously damaged, further leading to fibrosis formation, and eventually, esophageal stricture occurs. Therefore, early intervention of barrier dysfunction by improving esophageal barrier function may be the key to stopping the occurrence or development of esophageal strictures (Blevins *et al*, 2018; Kleuskens *et al*, 2022; Akdis, 2021).

Tacrolimus, or FK506, is a macrolide antibiotic and a powerful immunosuppressive agent. It binds to a cytosolic protein (FKBP12) to form the FKBP12 tacrolimus complex, which inhibits the calcium ion-dependent signaling pathway and further inhibits the release of interleukin-2 (IL-2) and the function of T lymphocytes(Wallemacq & Reding, 1993; Venkataramanan *et al*, 1995; Letko *et al*, 1999). In the clinic, FK506 plays a positive role in treating autoimmune diseases such as atopic dermatitis (AD), systemic lupus erythematosus (SLE), and autoimmune ophthalmopathy(Cury Martins *et al*, 2015; Fereig *et al*, 2021; Undre *et al*, 2009). Since FK506 is a medicine as an immunosuppressant for atopic dermatitis, it is applied to test the effects in solving inflammation and protecting the keratinocyte nature, where the treatment TA cannot work. This exploration has never been reported in the literature.

In this work, we first analyzed the biopsies of clinical esophageal stricture epithelial tissue (ESE) using proteomics technology. Transcriptomic information about the strictures was obtained from the gene expression omnibus (GEO) database to collect the relevant differentially expressed genes (DEGs). Cystatin A (CSTA) was thus identified. After then, the CSTA-mediated biological behaviors of human keratinocytes were explored. Gene CSTA in the cells was knocked down, followed by treatment of TA or tacrolimus (FK506). The keratinocyte behaviors were examined by scanning electron microscope (SEM), transmission electron microscopy (TEM), quantitative polymerase chain reaction (qPCR), immunofluorescence (IF), and flow cytometry (FC), as well as the effects of both drugs, TA and FK506, on the cells. The results showed that cell-cell distance enlargement, as well as cell-cell tight junctions, decreased after CSTA knockdown (kCSTA). Still, TA did not significantly protect kCSTA cells under the inflammatory environment. Unexpectedly, after FK506 treatment, the expression of CSTA, keratin 14(K14), cadherin 1 (CDH1), Zona occludens 1 (Zo-1), and β-Catenin in the cells increased significantly, suggesting that the kCSTA cells were rescued to some extent. It is supposed that FK506 would be a promising therapeutic medicine for patients with esophageal strictures in the presence of CSTA mutations, which increases cell-cell tight junctions and decreases cell-cell distances, restores esophageal epithelial barrier function, and thus stops the onset and progression of esophageal strictures as soon as possible. The workflow is displayed in **Figure 1**.

**Figure 1.**
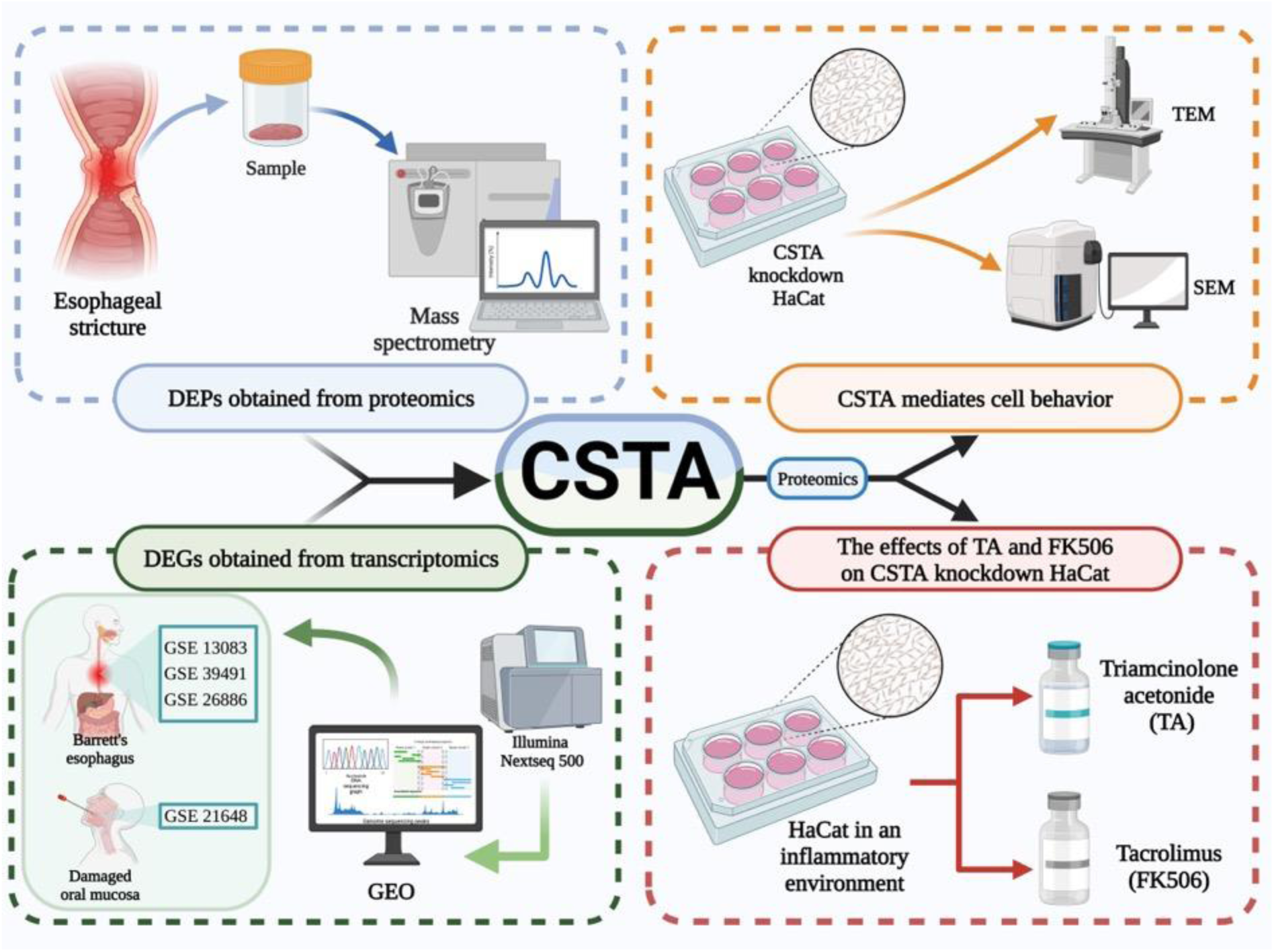
The workflow of this study.

## 2. Results

### 2.1 Proteomic Characteristics of Esophageal Stricture Epithelium

The disturbance of proteins in the esophageal stricture epithelial tissue (ESE) was investigated with LC-MS/MS analysis, as shown in **Figure 2**(a). The differential proteins of 172 up-regulated and 57 down-regulated were revealed in the esophageal stricture epithelial samples, compared with the normal esophageal epithelial tissue (NEE) (Figure 2(b-c)). The total differentially expressed proteins (DEPs) were analyzed through Gene Ontology (GO) and Kyoto Encyclopedia of Genes and Genomes (KEGG) pathway-enrichment analysis (Figure 2(d)). Furthermore, an interacting network among the top 50 DEPs consisting of co-expression, physical interaction, co-localization, predicted genetic interactions and shared protein domains was constructed through GeneMANIA (Figure 2(e)).

**Figure 2.**
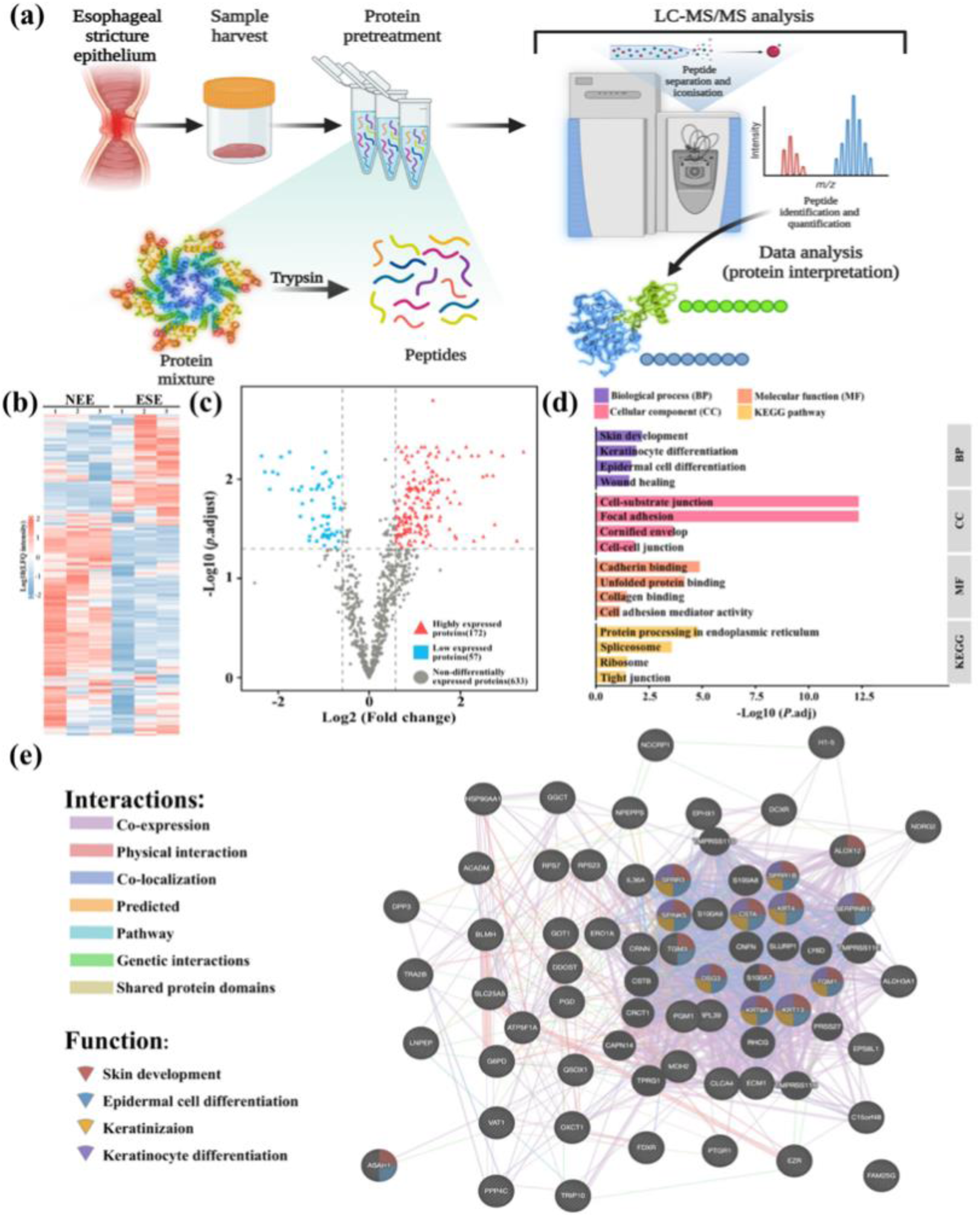
Proteomic analysis of esophageal stricture biopsy. (a) Proteomics protocol. (b) Heat map of differentially expressed proteins (DEPs). (c) Volcano map of proteomic analysis. (d) GO and KEGG pathway enrichment. (e) Protein-protein interacting networks among the top 50 DEPs.

The biological processes from GO analysis involved keratinocyte differentiation, epidermal cell differentiation, and wound healing. The KEGG pathway enrichment revealed that the tight junction between cells was enriched. Some specific proteins, such as CSTA, desmoglein 3 (DSG3), and K14, also enriched skin development, epidermal cell differentiation, keratinization, and keratinocyte differentiation, as shown in the protein-protein interaction network (Figure 2(e)). Because of the GO/KEGG enrichment results and the PPI results, we selected human keratin-forming cell lines (HaCaT), which are also widely distributed in long luminal organs such as the esophagus and urethra, for study in subsequent cell experiments. The proteomic results of epithelial samples from esophageal strictures predicted that esophageal epithelial barrier dysfunction occurs when esophageal strictures occur. The differentiation of epithelial cells, keratinocytes, and cell-cell junction plays a crucial role in the esophageal epithelial barrier (EEB).

Notably, the differentiation of keratinocytes and cell-cell junction gives an excellent barrier against harmful substances in foods or gastric acid refluxing from the stomach to protect the inner tissue of the esophagus. Under some conditions like pathological (e.g., Barrett’s esophagus) and physical injury (e.g., EMR/ESD), the EEB cannot function properly; the harmful substances containing antigens or/and hydrogen ions constantly attack basal cells to produce inflammation. When inflammation is not cured in time, it will stimulate fibroblasts to differentiate excessively into myofibroblasts, manifesting as excessive fibrosis and eventually leading to esophageal stricture (Blevins *et al*, 2018; Bouwstra *et al*, 2021). Therefore, timely reconstruction and repair of the esophageal epithelial barrier through treatment to enable the barrier to function correctly can fundamentally stop the occurrence and progression of esophageal strictures.

### 2.2 Screening and Obtaining the Distinctive Gene- CSTA

Barrett’s esophagus was selected as a representative disease of pathological stricture. Three datasets of Barrett’s esophagus, GSE13083, GSE26886, and GSE39491, were collected and analyzed to obtain co-upregulated, co-downregulated differentially expressed genes (DEGs) (**Figure S1**(a-b), Supporting Information). As we know, the oral mucosa is a representative tissue with perfect regenerative capability. Thereby, we retrieved the transcriptomics data of the oral mucosa and the cells from the NCBI Gene Expression Omnibus (GEO) database available at https://www##ncbi##nlm##nih##gov/geo/. For oral mucosal repair, 1 dataset, GSE21648, was searched and analyzed to obtain up-regulated and down- regulated DEGs (Figure S1(c), Supporting Information). Two Venn diagrams were obtained from the genes of up-regulated Barrett’s esophagus, down-regulated oral mucosa, and down-regulated Barrett’s esophagus and up-regulated oral mucosa (Figure S1(d), Supporting Information). The overlapped DEGs are shown in Table 1.

**Table 1.**
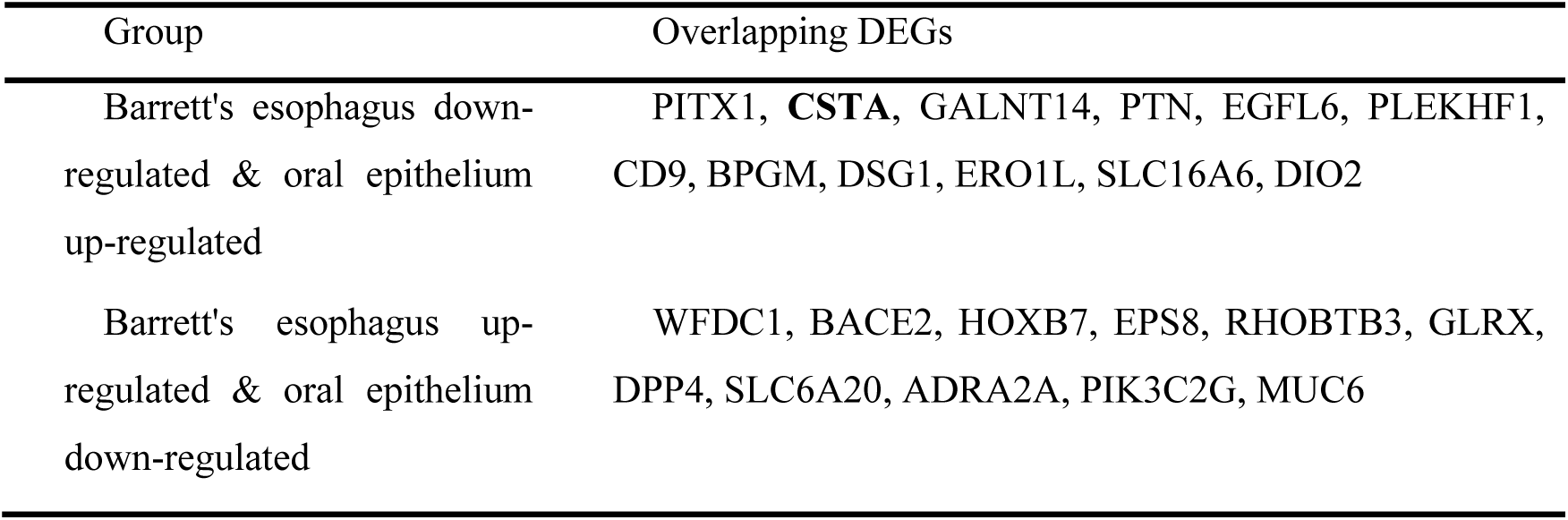
The overlapping DEGs from the DEGs of Barrett’s esophagus and oral epithelium.

| Group | Overlapping DEGs |
| --- | --- |
| Barrett's esophagus down-regulated & oral epithelium up-regulated | PITX1, **CSTA**, GALNT14, PTN, EGFL6, PLEKHF1, CD9, BPGM, DSG1, ERO1L, SLC16A6, DIO2 |
| Barrett's esophagus up-regulated & oral epithelium down-regulated | WFDC1, BACE2, HOXB7, EPS8, RHOBTB3, GLRX, DPP4, SLC6A20, ADRA2A, PIK3C2G, MUC6 |

PPI analysis of these two overlapping DEGs suggested that these DEGs are mainly on epithelial cell differentiation and keratin formation (**Figure S2,** Supporting Information). This result is the same as that of the PPI analysis of proteomics (Figure 2(e)).

Overlapping these DEGs with the DEPs taken from the proteomics of esophageal stricture biopsy, CSTA was discovered. It was downregulated in DEPs of esophageal strictures (the proteomics, Figure 2(c), and downregulated in DEGs of Barrett’s esophagus but upregulated in DEGs of the oral mucosa (the transcriptomics, Table 1)). In previous studies, CSTA was found to play an essential role in maintaining the epithelial barrier and epithelial cell development (Candi *et al*, 2005; Butler *et al*, 2011; Pernu *et al*, 1990). It has also been reported to play a role in maintaining the morphology of the cornified envelope as a precursor protein and was proposed as a diagnostic tool for head, neck, and chest cancer and prognosis (Blaydon *et al*, 2011; Moosbrugger-Martinz *et al*, 2015; Wei *et al*, 2022; Gupta *et al*, 2015; Candi *et al*, 2005). Therefore, CSTA can be proposed as a potential biomarker for esophageal strictures, which has never been reported in the literature.

In the stricture esophagus, CSTA mutated and resulted in the weak function of EEB, which allowed harmful substances to attack basal cells and further produced constant inflammation. Many inflammatory factors were secreted and continuously stimulated the conversion of fibroblasts in the connective tissue into myofibroblasts, leading to esophageal fibrosis and further stricture. The mutation of CSTA might be the cause of esophageal strictures in some patients. That might be why patients with esophageal strictures are not sensitive to the therapies with glucocorticoid medicines.

### 2.3 The Relationship Between Knockdown CSTA Cells and Epithelial Barrier Dysfunction

First, we verified the protein levels of CSTA and ZO-1 in the epithelium of esophageal strictures by IHC images in the human protein atlas (HPA) database and WB to better combine the results obtained from the above proteomics (IHC images in the HPA database were used because the single biopsy sample was too small for immunohistochemical staining due to surgical manipulation and ethical restrictions, and WB was chosen for verification of protein expression levels). By IHC, CSTA and ZO-1 were found to be expressed in the normal esophagus. By detecting protein levels by WB, we found that CSTA and ZO-1 protein levels were higher in NEE than in ESE, validating the above proteomics results for CSTA. ZO-1 protein levels indicated weakened cell-cell tight junctions in the esophageal stricture epithelium, suggesting dysfunction of the EEB (**Figure 3**(a-b)).

**Figure 3.**
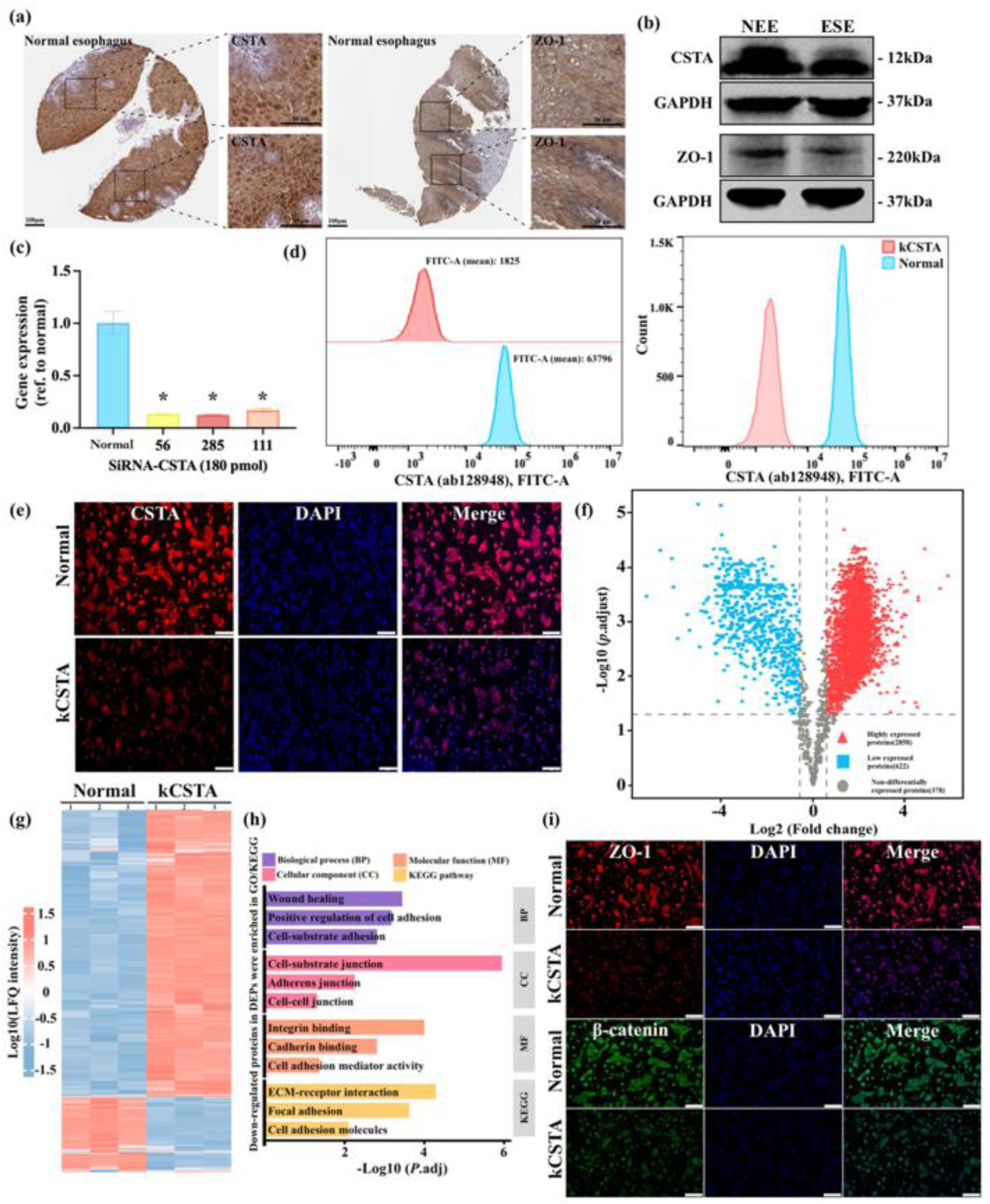
The relationship between knockdown CSTA cells and epithelial barrier dysfunction. (a) Immunohistochemical (IHC) staining for CSTA and ZO-1 in normal esophageal epithelium from the HPA database. Scale bars, 100 μm (left panels), 50 μm (right panels). (b) CSTA and ZO-1 protein expression in biopsy samples were analyzed by Western blot. (c) Expression of CSTA at 180 pmol for different base sequences. Cells were transfected by SiRNA-CSTA for 24 h and tested with qPCR technology. (d) Expression of CSTA after transfected by SiRNA-CSTA for 24 h and analyzed with flow cytometry assay. (e) Immunofluorescent staining with anti-CSTA as the primary antibody. \**P*< 0.05, compared with the normal HaCaTs. Scale bars, 200 μm. (f) Volcano plot of differentially expressed proteins in knockdown CSTA cells versus normal cells, red represents high expression, blue represents low expression, and gray represents no significant difference. (g) Heat map of differentially expressed proteins in knockdown CSTA cells (kCSTA) versus normal cells (Normal). (h) GO/KEGG pathway enrichment results for down-regulated proteins in DEPs after knockdown of CSTA expression. (i) Immunofluorescence staining for ZO-1 and β-catenin in kCSTA cells after knockdown of CSTA expression.

Since epithelial cells are the most dominant cells performing EEB functions, keratinocytes are widely present in cavity-like organs in the body. As demonstrated by the proteomic results of epithelial samples from esophageal strictures, keratinocytes were selected to test their subsequent biological behavior and sensitivity to clinical medicine.

The function of CSTA in esophageal stricture has never been reported in previous studies. Herein, we explored the role of CSTA in esophageal keratinocytes with HaCaT as the model cells. The gene CSTA in HaCaT cells was knocked down using siRNA technology as kCSTA cells. The knockdown efficiency, up to more than 80%, was verified by qPCR assay compared with the normal cells (Figure 3(c)). The expression of CSTA was significantly reduced in the kCSTA cells. Similar results were observed in flow cytometry assay and immunofluorescent analysis, where the protein expression of CSTA significantly reduced in the kCSTA cells with the normal cells as the reference (Figure 3(d-e)).

To further clarify the role of CSTA in cells, we performed proteomic analysis of kCSTA cells using normal cells as a control to obtain DEPs after knocked down of CSTA expression. The differential proteins of 2858 up-regulated and 622 down-regulated were revealed in the kCSTA cells, compared with the normal cells (Figure 3(f-g)). The up-regulated and down-regulated proteins were analyzed separately through GO and KEGG pathway-enrichment analysis (Figure 3(h) and **Figure S4,** Supporting Information). The down-regulated proteins were mainly enriched in cell adhesion and cell junction-related pathways, such as positive regulation of cell adhesion, cell-cell junction, cell adhesion mediator activity, etc. Such results suggest that the knockdown of CSTA expression results in decreased cell-cell adhesion and cell junction. Cell-cell adhesion and cell junctions were reduced after knocked down of CSTA expression. We then performed immunofluorescence staining of kCSTA cells. We found that the levels of ZO-1 and β-Catenin, those proteins representing cell-cell tight junctions, were reduced after knocked down of CSTA expression (Figure 3(i)). This further validated the above proteomic results.

Then, to verify the above results, we again observed the behavior of cells after knocking down CSTA expression under SEM and TEM. HaCaT cultured under normal conditions displayed cobblestone morphology. Cells assembled and adhered with many junctions between others (**Figure 4**(a)). After CSTA was knocked down, cells scattered from each other, and large gaps between cells appeared. Similar results were observed in TEM measurement (Figure 4(b)). This result indicated that the reduced CSTA in cells significantly inhibited the cell junction, thereby enlarging the distance between cells.

**Figure 4.**
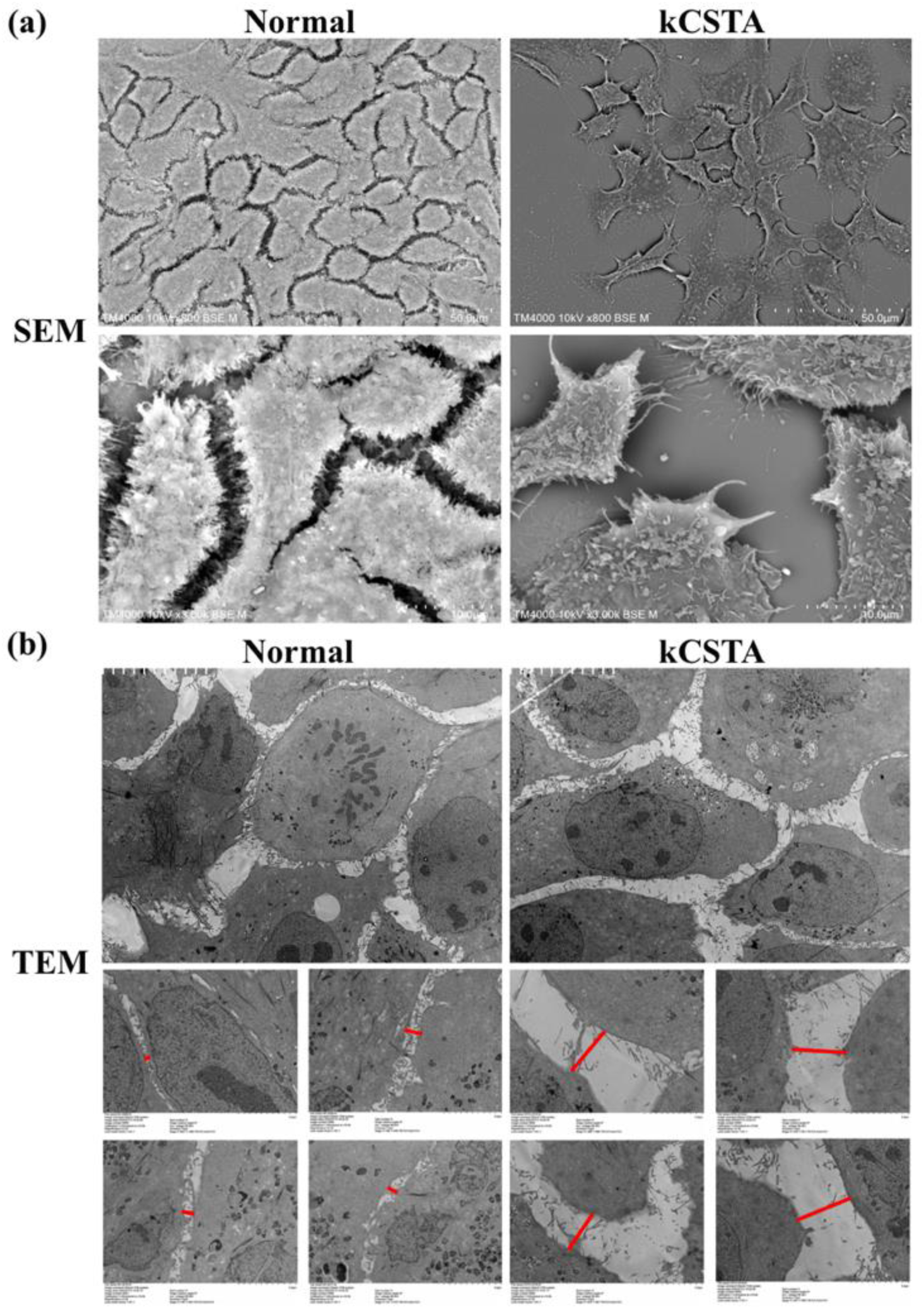
Cell-cell junctions were observed under SEM (a) and TEM (b). Cells were treated to be CSTA knockdown (kCSTA), with normal cells as the reference. The red lines denote the distance between cells.

Gene CSTA encodes protein Cystatin A. It is a protease inhibitor and is one of the precursor proteins of cornified cell envelope in keratinocytes and plays a role in epidermal development and maintenance. It is also an intracellular inhibitor regulating the activities of cystatin proteinase and has a vital role in desmosome-mediated cell-cell adhesion (Blaydon *et al*, 2011; Candi *et al*, 2005). In our work, gene CSTA was highly correlated with the tight junctions between HaCaT cells. The intercellular adhesion was abnormally weakened, and the intercellular gap became abnormally large when CSTA expression was suppressed.

In esophageal strictures, the gap between the epithelial cells increases, resulting in a dysfunctional EEB. The basal cells are more susceptible to hydrogen ion refluxing from the stomach or/and some harmful antigens from foods, producing persistent inflammation and, consequently, fibrosis and finally leading to esophageal strictures. This suggests that the normal expression of CSTA is necessary for the proper function of the EEB. After ESD/EMR, some patients with CSTA mutations may need to mediate normal CSTA expression to promote good EEB function and stop esophageal strictures’ onset and progression.

### 2.4 Effect of Triamcinolone Acetonide (TA) on kCSTA Cells

TA is a glucocorticoid medicine used in the clinic for anti-inflammation with local injections or/and oral therapy. It was employed to treat kCSTA cells to simulate the clinical state. HaCat cells were cultured for 24 h in an inflammatory environment (supplementary with IL-1β and TNF-α, as literature (Blaydon *et al*, 2011; Blawat *et al*, 2020; Zhao *et al*, 2020) reported). The expressions of keratin 14 (K14), cadherin 1 (CDH1), Zona occludens 1 (ZO-1), and β-Catenin were tested. K14 is expressed explicitly in epidermal keratinocytes and is often used as a biomarker in keratinocyte-driven re-epithelialization (Candi *et al*, 2005; Meyer *et al*, 2014; Wang *et al*, 2021; Korbut *et al*, 2020). CDH1, encoded by gene CDH1, is a calcium-dependent cell adhesion protein (Candi *et al*, 2005; van Roy & Berx, 2008; Zhang *et al*, 2019). It plays a critical role in cell-cell adhesion and tissue organization. ZO-1 is a tight junction (TJ) protein found in complexes at cell-cell contacts. The role of ZO-1 is to recruit other TJ proteins. The protein is localized to the cytoplasmic membrane, mainly where it is essential for barrier function(Kuo *et al*, 2021; Dobson *et al*, 1994). β-catenin, or CTNNB1, is an evolutionarily conserved, multifunctional intracellular protein. β-catenin was originally identified in cell adhesion junctions (AJs), where it functions to link the cytoplasmic domains of cadherins to the a-catenin and actin cytoskeletons in response to cell-cell adhesion(Nakayama *et al*, 2014; Puthia *et al*, 2014; Wang *et al*, 2022). CDH1 and β- catenin form a complex in cells. This complex is a critical component of the cell-cell adhesion complex(Wang *et al*, 2022; Liu *et al*, 2022; Jovov *et al*, 2011). Thus, in this work, K14 was measured as a biomarker of HaCaT cells, while CDH1, β-catenin, and ZO-1 were measured as functional biomarkers of cell-cell adhesion.

The cell growth was significantly reduced after they were cultured in an inflammatory environment. The intercellular adhesion weakened (**Figure 5**(a-d)). The expression of K14, CDH1, β-catenin, and ZO- 1 decreased obviously in the IE group than in the normal group, indicating that the cells lived abnormally in an inflammatory condition (Figure 5(a-d), fluorescent intensity). Interestingly, the cells grew and proliferated after TA treated them for 24 h. The cell number and fluorescent intensity increased, compared with those in the IE group (Figure 5(a-d), IE/TA), indicating that TA can protect cells from attack by the inflammatory environment. The results of the CCK8 assay shows enhanced cell proliferation with TA in the inflammatory environment (IE/TA) compared to the IE group (**Figure S3**, Supporting Information). However, after CSTA was knocked down, many cells (kCSTA/IE/TA group) died. And the expressions of K14, CDH1, β-catenin, and ZO-1 reduced significantly compared with those of normal and IE/TA cells. TA cannot produce a normal effect on kCSTA cells, an effect that keeps the cells from being attacked by the inflammatory environment.

**Figure 5.**
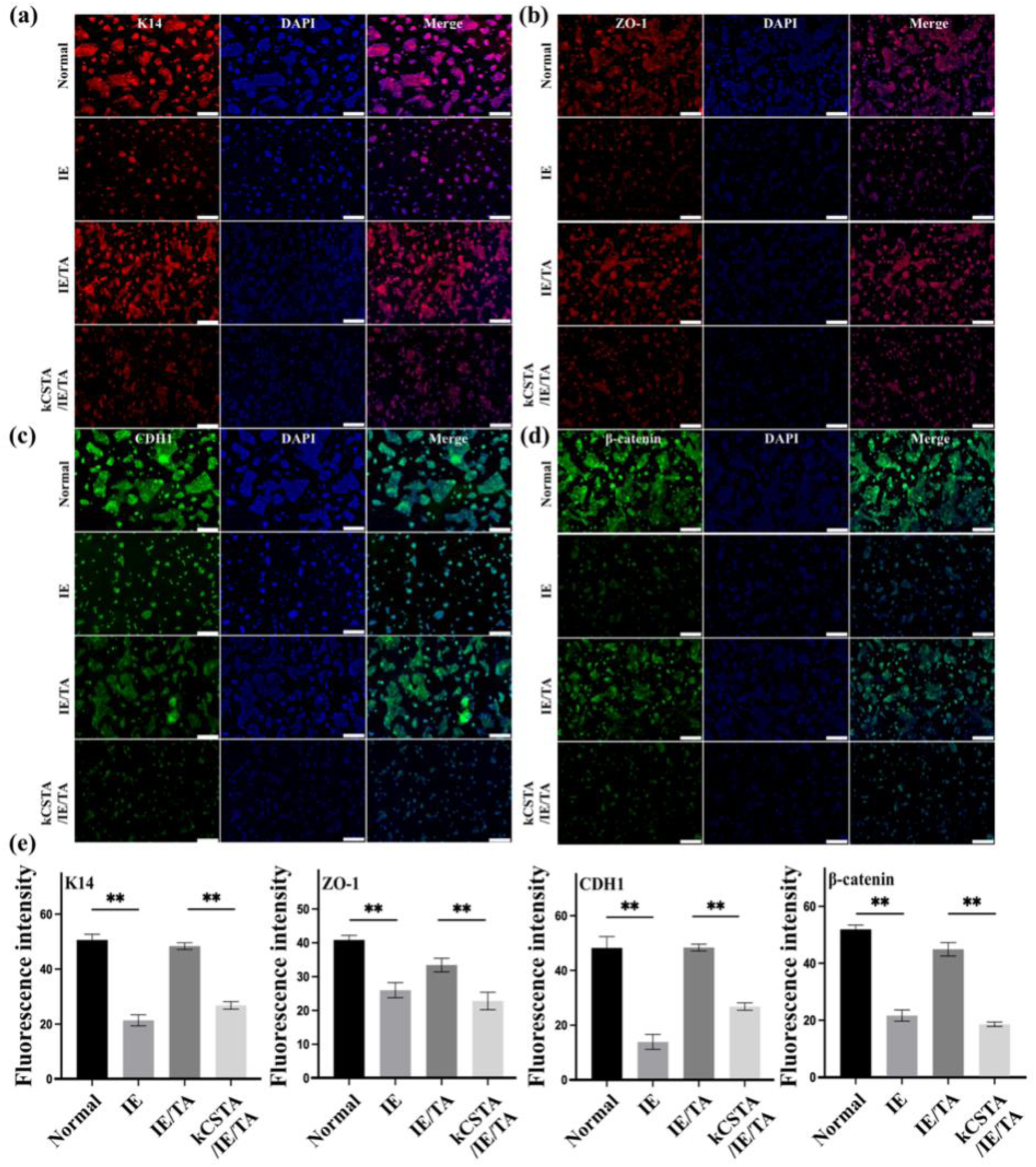
Construction of a clinical inflammatory environment and the effect of TA on cells. Construction of a clinical inflammatory environment and the effect of TA on cells. Cells were cultured for 24 h in the medium supplementary with TNF-α (10 ng mL^-1^) and IL-1β (5 ng mL^-1^). Normal cells were cultured in the inflammatory medium, simplified as the IE group. Normal cells and kCSTA cells were cultured in an inflammatory environment and treated by TA, named IE/TA and kCSTA/IE/TA, respectively. (a) Immunofluorescent (IF) staining with anti-K14 as the primary antibody. (b) IF staining with anti-ZO-1 as the primary antibody. (c) IF staining with anti-CDH1 as the primary antibody. (d) IF staining with anti-β-catenin as the primary antibody. (e) Quantitative analysis of the above immunofluorescence by Image J.. \**P*< 0.05, \*\**p*<0.01, compared with the normal cell. Scale bars, 200 μm.

The high expressions of K14, CDH1, β-catenin, and ZO-1 in the IE/TA group confirmed the therapeutic effect of TA on cells in an inflammatory state, which is similar to some patients with esophageal strictures using the local injection of TA in the surgical area before ESD/EMR (Nagami *et al*, 2018; Kato *et al*, 2018; Wang *et al*, 2023; Sakaguchi *et al*, 2020). In the effect of TA, CSTA-unmutated epithelial cells can resist the effects of the inflammatory environment on cells. Cells exhibit normal intercellular adhesion and tight junctions, thus restoring EEB function, and EEB function is fixed within the effective time of glucocorticoids, allowing harmful substances to attack the underlying tissue as little as possible and reducing the inflammatory response, suggesting that glucocorticoids can relieve esophageal strictures and produce a therapeutic effect.

However, TA does not work for kCSTA cells in the inflammatory microenvironment; it does not make the cells resistant to the inflammatory environment and does not elevate the expression of K14, CDH1, β-catenin, and ZO-1. This explains why a proportion of patients with clinical esophageal strictures fail to relieve after glucocorticoid treatment. The mutation of CSTA weakens the adhesion and tight junctions between epithelial cells, leading to EEB dysfunction. During the effective time of glucocorticoids, the function of EEB does not improve due to the mutation of CSTA, and harmful substances continue to invade the underlying tissues, leading to continuous inflammation and eventually fibrosis, i.e., esophageal stricture. Therefore, some patients are insensitive to glucocorticoids and still develop esophageal strictures.

### 2.5 Effect of Tacrolimus (FK506) on kCSTA Cells

Tacrolimus, also named FK506, is an immunosuppressive medicine. It is often applied to treat atopic dermatitis but never treated strictures before. We chose it as a candidate medicine for treating kCSTA cells where the TA did not work.

By immunofluorescence assay, the expressions of protein K14, CDH1, ZO-1, and β-catenin were significantly higher in the FK506-treated cells (kCSTA/IE/FK506) than in the TA-treated cells (kCSTA/IE/TA), both in the inflammatory environment. There was no statistically significant difference in protein expressions between the control (kCSTA/IE) and kCSTA/IE/TA (**Figure 6**(a-e)). By flow cytometry assay, the expressions of both K14, CDH1, ZO-1, and β-catenin were also higher in the FK506-treated cells than in the TA-treated or the control cells (Figure 6(f)). The results of the CCK8 assay shows enhanced cell proliferation with FK506 compared to TA (Figure S3, Supporting Information). These results suggest that FK506 has a better effect on CSTA-knockdown cells in the inflammatory environment than TA.

**Figure 6.**
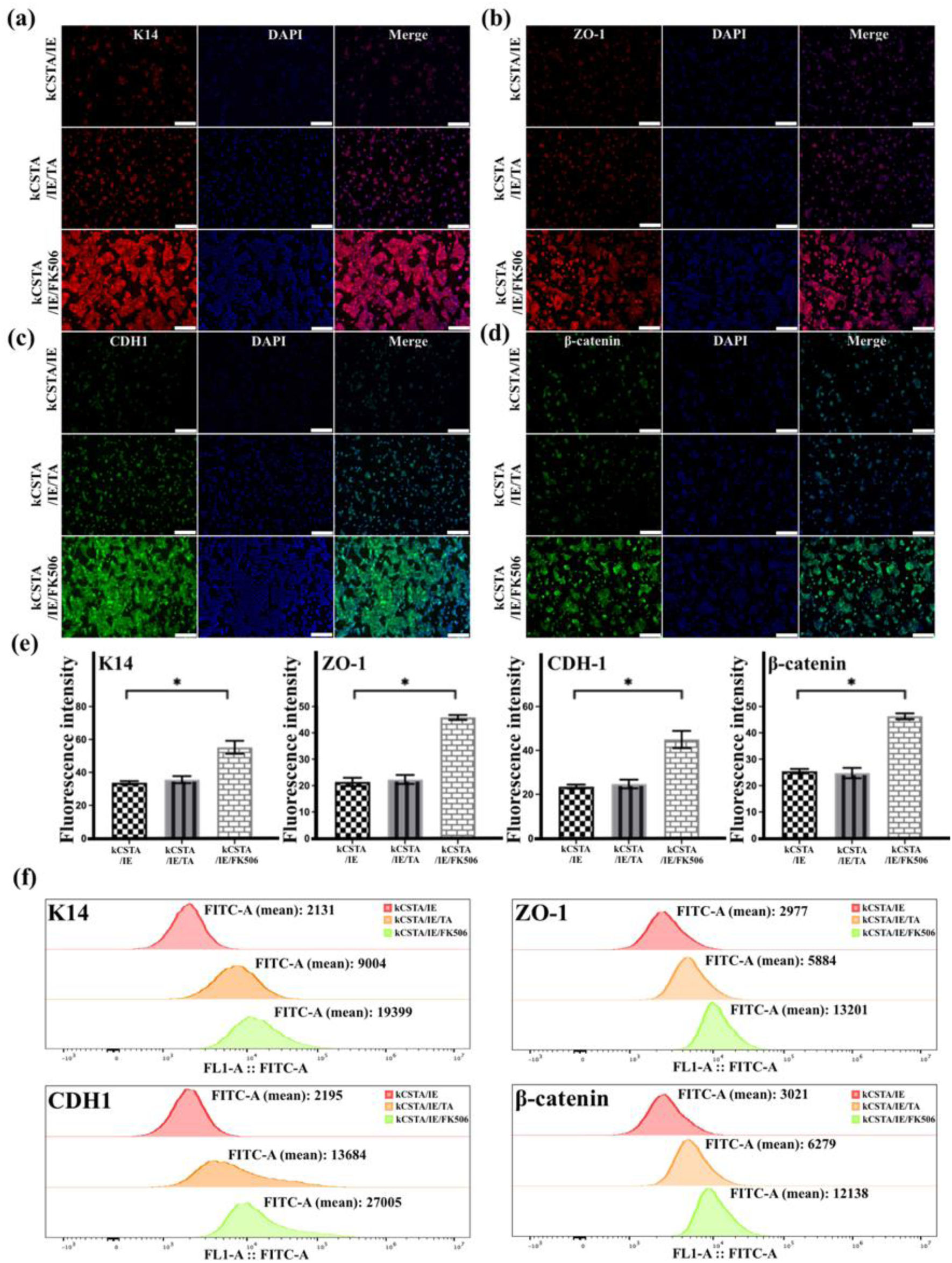
Effect of TA or FK506 on kCSTA cells in an inflammatory environment. kCSTA cells were cultured in an inflammatory environment called kCSTA/IE cells. The protein expressions of K14, CDH1, ZO-1, and β-catenin were tested after cells were treated for 24 h by TA or FK506, respectively. (a) IF staining with anti-K14 as the primary antibody. (b) IF staining with anti-ZO-1 as the primary antibody. (c) IF staining with anti-CDH1 as the primary antibody. (d) IF staining with anti-β-catenin as the primary antibody. (e) Quantitative analysis of the above immunofluorescence by Image J. (f) Flow cytometry assay. \**P*< 0.05, compared with the kCSTA/IE. Scale bars, 200 μm.

In previous studies, some researchers have found that CSTA is highly associated with atopic dermatitis. Some immunosuppressive medicines are employed to treat atopic dermatitis, with tacrolimus (FK506) being particularly effective among them (Cury Martins *et al*, 2015; Fereig *et al*, 2021; Undre *et al*, 2009). However, immunosuppressive medication has not been used in patients with esophageal strictures, and no relevant literature has been reported.

Tacrolimus (FK506), an immunosuppressive medicine, is widely used in clinical treatments for diseases like immune rejection after organ transplantation, atopic dermatitis, and T-cell-mediated diseases (Fereig *et al*, 2021; Undre *et al*, 2009; Wallemacq & Reding, 1993). Recent studies have shown that FK506 has a therapeutic effect on inflammation caused by eosinophilia (Kandikattu *et al*, 2021). A proportion of patients with esophageal strictures are due to eosinophilic esophagitis. Thus, we conjecture that FK506 will take function for those patients. That’s why FK506 was used as the candidate to treat kCSTA cells. The above results confirmed elevated growth of kCSTA cells treated with FK506 but not with TA; expression of K14, CDH1, ZO-1, and β-catenin was increased. This result implies that the use of FK506 in CSTA mutant epithelial cells in an inflammatory state causes the cells to exhibit excellent status and relatively tight cell-cell junctions, allowing the maintenance of EEB function, protection from harmful substances, and a safe growth environment for basal cells. FK506 restores the dysregulated EEB function (mutation of CSTA) and resists the effects of the inflammatory environment on the cells, allowing epithelial cells to restore function while reducing the inflammatory response. This predicts that FK506 is a promising agent for resisting the development and progression of esophageal strictures.

### 2.6 CSTA Expression of Cells Treated by TA or FK506

To verify the hypothesis that CSTA is closely associated with esophageal stricture epithelium and might be a novel target of immunosuppressive medicine, the expression of CSTA was investigated after kCSTA cells were treated by TA or FK506. The results showed that CSTA expressions in both kCSTA/IE or kCSTA/IE/TA were much lower than that in the normal cells, though the expression in the kCSTA/IE/TA group was a little higher than that in kCSTA/IE group. However, cells treated with FK506 (kCSTA/IE/FK506) expressed more CSTA proteins than kCSTA/IE/TA group (**Figure 7**(a)), indicating that FK506 is an effective therapy for cells with kCSTA in inflammatory conditions. The results also showed that both TA- or FK506-treated cells expressed less CSTA than the normal cells, which might be attributed to the short time or low frequency of medicine administration.

**Figure 7.**
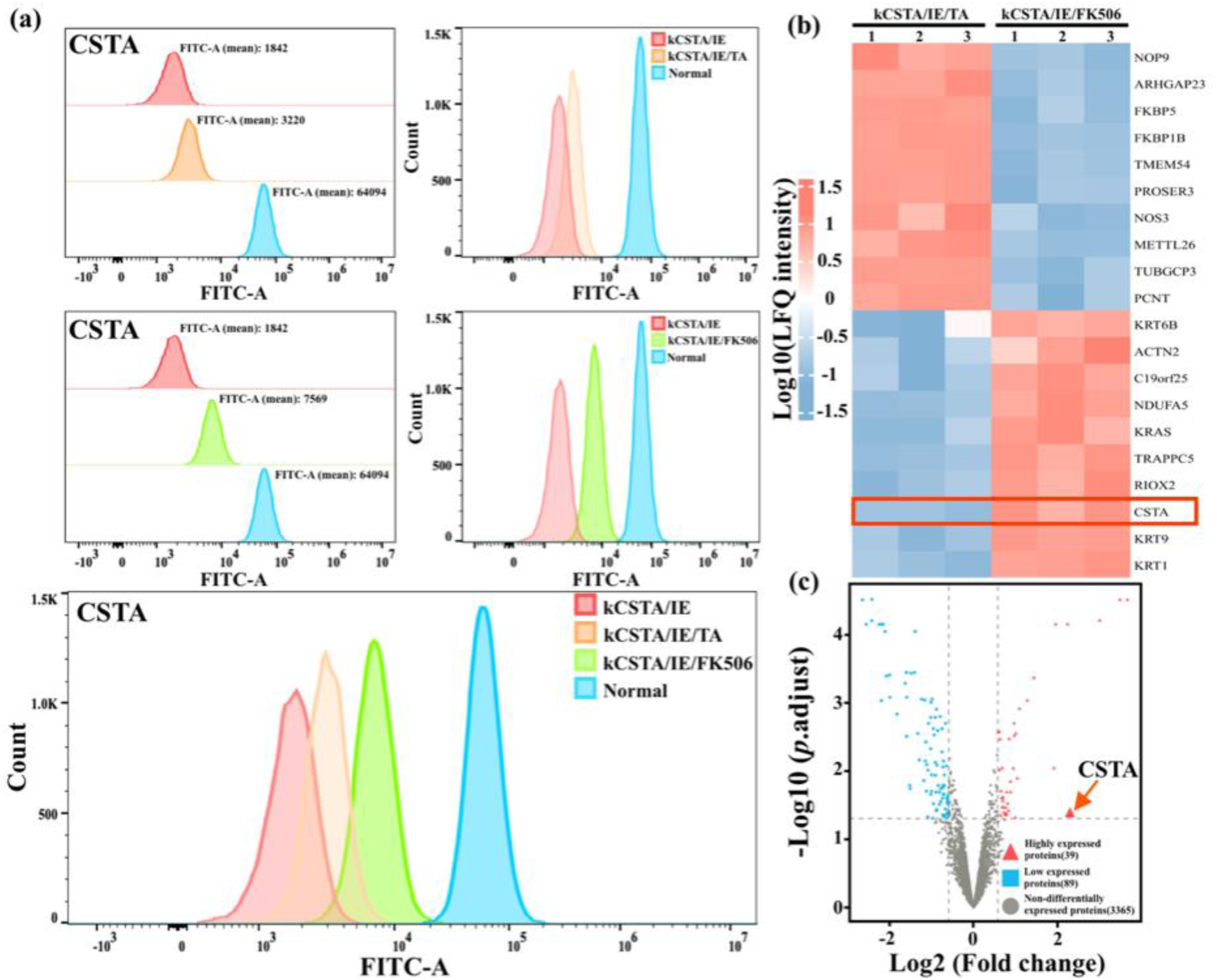
Effects of TA and FK506 on CSTA expression in kCSTA cells using flow cytometry assay and proteomics. (a) Analysis of CSTA protein expression by flow cytometry. The Cells with CSTA knockdown was cultured for 24 h in the inflammatory condition (simplified as kCSTA/IE), followed by TA (kCSTA/IE/TA) or FK506 (kCSTA/IE/FK506) treatment, with the normal cells under normal culture conditions as the positive control. (b) Heatmap of the top 10 up- and down-regulated DEPs in the proteomics of cells after being treated with TA and FK506, respectively. (c) Volcano plots of DEPs in proteomics of cells treated with TA and FK506, respectively.

After this, we again performed proteomic analysis using mass spectrometry on FK506-treated kCSTA cells (kCSTA/IE/FK506) and TA-treated kCSTA cells (kCSTA/IE/TA), respectively, to further validate the above point. The differential proteins of 39 up-regulated and 89 down-regulated were revealed in the kCSTA/IE/FK506, compared with the kCSTA/IE/TA. The top 10 most differentially expressed EDPs (up- and down-regulated) are shown in a heat map, as demonstrated by the flow cytometric analysis above, and by proteomics, we also found that CSTA was significantly more highly expressed in FK506- treated kCSTA cells, compared to TA-treated kCSTA cell (Figure 7(b-c)).

In previous experiments, we found that TA could positively affect cells without CSTA knockdown but had a poor effect on kCSTA cells. This may be related to the inability of TA itself to enhance CSTA expression in cells, which in turn fails to restore cell-cell adhesion and tight junctions, resulting in failure to restore EEB function. In this case, FK506 could be used, which elevated the expression of CSTA and promoted the expression of K14, CDH1, ZO-1, and β-catenin. This suggests that epithelial cells are aided by the drug FK506 both to restore EEB function by enhancing cell-cell tight junctions and adhesion and to resist the effects of the inflammatory environment on the cells. Therefore, we believe patients with stricture with CSTA mutations and insensitivity to glucocorticoid treatment can be treated with FK506. Alternatively, if the patient is unsure whether the CSTA is mutated in the epithelium, the new idea is to mix FK506 with TA (similar to cocktail therapy). Indeed, this is a new therapy or hope for treating esophageal strictures. However, more testing or clinical trials are needed before treating patients.

## 3. Discussion

Following esophageal ESD/EMR, the surgical damage and inflammation within the microenvironment impact the functioning of the epithelial cells(Hsu *et al*, 2023; Kanai *et al*, 2012). This leads to reduced cell-cell adhesion and tight junctions, resulting in a dysfunction of the epithelial barrier(Li *et al*, 2019; Kiesslich *et al*, 2012; Zhang *et al*, 2021). Consequently, harmful substances persistently attack the underlying tissues, triggering chronic inflammation that perpetuates fibroblast over-differentiation and eventually causes esophageal stricture. Therefore, dysfunction of the esophageal barrier plays a crucial role in initiating esophageal strictures. Swift restoration of the esophageal epithelial barrier function can effectively prevent and alleviate the progression of esophageal stricture. Glucocorticoids have emerged as a highly effective approach for treating or alleviating esophageal strictures. While some patients experience relief from esophageal stricture with glucocorticoids, others may exhibit resistance to their effects, and the underlying cause for this insensitivity remains unclear (Hsu *et al*, 2023; Sakaguchi *et al*, 2023; Koizumi *et al*, 2023; Hikichi *et al*, 2020).

Here, we first obtained the DEPs by proteomic analysis of biopsy tissues from patients with esophageal strictures. Then, the data sets of esophageal stricture disease represented by Barrett’s esophagus and oral mucosa were collected separately from the GEO database to obtain the DEGs for the corresponding intersection. Comparing DEPs with DEGs, we identified CSTA as a potential biomarker for esophageal stricture disease. Subsequently, we explored the CSTA-mediated cell biological behavior and performed proteomic analysis of cells after the knockdown of CSTA. The decreased expression of ZO-1 and β-catenin proteins and the expanded cell gaps under SEM and TEM confirmed the diminished cell-cell tight junctions and adhesion after the knockdown of CSTA. After mimicking the inflammatory environment, the drug TA could not make kCSTA cells resist the inflammatory environment and showed TA insensitivity; the expression of K14, CDH1, ZO-1, and β-catenin significantly decreased. However, after FK506 treatment, kCSTA cells exhibited higher protein expression than after TA treatment.

Moreover, we also verified by proteomic analysis that a significantly higher expression of CSTA occurs after the use of FK506 compared to the use of TA. The increased expression of K14, CDH1, ZO- 1, and β-catenin after the use of FK506 indicated that the influence of the inflammatory environment on kCSTA cells was alleviated, and the tight junctions and adhesion of keratinocytes increased, predicting rescue of dysregulated EEB function. Thus, CSTA may be a novel biomarker for restoring the epithelial barrier in esophageal strictures and could be a potential target for medical therapy.

One limitation of this study is the differential expression of CSTA between humans and mice (or other rodents), which hinders the validation of CSTA’s function through the creation of a mouse model for esophageal stricture. To address this limitation, future research endeavors will focus on validating the relationship between CSTA and the esophageal epithelial barrier in large animals that closely resemble humans. Additionally, more comprehensive investigations will be conducted in large animal experiments to explore detailed aspects such as the optimal timing and dosage of FK506 administration. These endeavors will provide a stronger theoretical foundation for conducting further clinical trials aimed at developing future clinical treatments.

Overall, CSTA plays a crucial role in maintaining the integrity of the esophageal epithelial barrier. A mutation in CSTA leads to a decrease in cell-cell adhesion and tight junctions, ultimately resulting in dysfunction of the esophageal epithelial barrier and the development of esophageal strictures. Furthermore, CSTA mutations also render individuals insensitive to glucocorticoid therapy. In this context, FK506 shows promise as a potential therapeutic option to restore esophageal epithelial function by enhancing CSTA expression. Additionally, combining FK506 with TA as a cocktail therapy could be a groundbreaking approach to inhibit the onset and progression of esophageal strictures, offering relief to affected individuals. This is the focus of our research group’s current efforts.

## 4. Methods

Matched clinical samples of esophageal stricture epithelium and adjacent normal esophageal tissue were obtained from three patients, provided by the Affiliated Hospital of Ningbo University, under written informed consent from patients before ESD/EMR surgery. The collection of human samples and the investigation protocols were approved by the Institutional Ethics Committee of Ningbo University (Ethical Committee approval No. KY20190101). Informed consent of all participating subjects was obtained. Detail Materials and methods are described in Supporting Information

## Supporting Information

Supporting Information is available from the https://www.embopress.org/journal/14602075 or from the author.

## Acknowledgements

We acknowledge the financial support from the National Natural Science Foundation of China (U20A20121), the One Health Interdisciplinary Research Project of Ningbo University (HY202210), and the Major Project of 2025 Sci & Tech Innovation of Ningbo (2020Z096). This work was also sponsored by K.C. Wang Magna/Education Fund of Ningbo University. Thanks for the technical support by the Core Facilities, Health Science Center of Ningbo University.

## Conflict of interest

All other authors declare they have no competing interests.

Received: ((will be filled in by the editorial staff))

Revised: ((will be filled in by the editorial staff))

Published online: ((will be filled in by the editorial staff))

## References

1. Akdis CA (2021) Does the epithelial barrier hypothesis explain the increase in allergy, autoimmunity and other chronic conditions? Nat Rev Immunol 21: 739–751

2. Blawat K, Mayr A, Hardt M, Kirschneck C, Nokhbehsaim M, Behl C, Deschner J, Jäger A & Memmert S (2020) Regulation of Autophagic Signaling by Mechanical Loading and Inflammation in Human PDL Fibroblasts. Int J Mol Sci 21: 9446

3. Blaydon DC, Nitoiu D, Eckl K-M, Cabral RM, Bland P, Hausser I, van Heel DA, Rajpopat S, Fischer J, Oji V, et al (2011) Mutations in CSTA, encoding Cystatin A, underlie exfoliative ichthyosis and reveal a role for this protease inhibitor in cell-cell adhesion. Am J Hum Genet 89: 564–571

4. Blevins CH, Iyer PG, Vela MF & Katzka DA (2018) The Esophageal Epithelial Barrier in Health and Disease. Clin Gastroenterol Hepatol 16: 608–617

5. Bouwstra JA, Helder RWJ & El Ghalbzouri A (2021) Human skin equivalents: Impaired barrier function in relation to the lipid and protein properties of the stratum corneum. Adv Drug Deliv Rev 175: 113802

6. Butler MW, Fukui T, Salit J, Shaykhiev R, Mezey JG, Hackett NR & Crystal RG (2011) Modulation of cystatin A expression in human airway epithelium related to genotype, smoking, COPD, and lung cancer. Cancer Res 71: 2572–2581

7. Candi E, Schmidt R & Melino G (2005) The cornified envelope: a model of cell death in the skin. Nat Rev Mol Cell Biol 6: 328–340

8. Cury Martins J, Martins C, Aoki V, Gois AFT, Ishii HA & da Silva EMK (2015) Topical tacrolimus for atopic dermatitis. Cochrane Database Syst Rev 2015: CD009864

9. Dobson H, Pignatelli M, Hopwood D & D’Arrigo C (1994) Cell adhesion molecules in oesophageal epithelium. Gut 35: 1343–1347

10. Duivenvoorden HM, Rautela J, Edgington-Mitchell LE, Spurling A, Greening DW, Nowell CJ, Molloy TJ, Robbins E, Brockwell NK, Lee CS, et al (2017) Myoepithelial cell-specific expression of stefin A as a suppressor of early breast cancer invasion. J Pathol 243: 496–509

11. El-Asmar KM, Hassan MA, Abdelkader HM & Hamza AF (2013) Topical mitomycin C application is effective in management of localized caustic esophageal stricture: a double-blinded, randomized, placebo-controlled trial. J Pediatr Surg 48: 1621–1627

12. Fereig SA, El-Zaafarany GM, Arafa MG & Abdel-Mottaleb MMA (2021) Tacrolimus-loaded chitosan nanoparticles for enhanced skin deposition and management of plaque psoriasis. Carbohydr Polym 268: 118238

13. Griffin MF, Fahy EJ, King M, Guardino N, Chen K, Abbas DB, Lavin CV, Diaz Deleon NM, Lorenz HP, Longaker MT, et al (2022) Understanding Scarring in the Oral Mucosa. Adv Wound Care (New Rochelle) 11: 537–547

14. Gupta A, Nitoiu D, Brennan-Crispi D, Addya S, Riobo NA, Kelsell DP & Mahoney MG (2015) Cell cycle- and cancer-associated gene networks activated by Dsg2: evidence of cystatin A deregulation and a potential role in cell-cell adhesion. PLoS One 10: e0120091

15. Hashimoto S, Kobayashi M, Takeuchi M, Sato Y, Narisawa R & Aoyagi Y (2011) The efficacy of endoscopic triamcinolone injection for the prevention of esophageal stricture after endoscopic submucosal dissection. Gastrointest Endosc 74: 1389–1393

16. Hikichi T, Nakamura J, Takasumi M, Hashimoto M, Kato T, Kobashi R, Takagi T, Suzuki R, Sugimoto M, Sato Y, et al (2020) Prevention of Stricture after Endoscopic Submucosal Dissection for Superficial Esophageal Cancer: A Review of the Literature. J Clin Med 10: 20

17. Horváth ÖP, Pavlovics G, Cseke L, Vereczkei A & Papp A (2023) Dysphagia After Esophageal Replacement and Its Treatment. Dysphagia

18. Hsu W-H, Shih H-Y, Shen C-S, Yu F-J, Wang H-C, Chan L-P, Kuo C-H, Hsieh H-M & Wu I-C (2023) Prevention and management of esophageal stricture after esophageal ESD: 10 years of experience in a single medical center. J Formos Med Assoc 122: 486–492

19. Iwata K, Mikami Y, Kato M, Yahagi N & Kanai T (2021) Pathogenesis and management of gastrointestinal inflammation and fibrosis: from inflammatory bowel diseases to endoscopic surgery. Inflamm Regen 41: 21

20. Jovov B, Que J, Tobey NA, Djukic Z, Hogan BLM & Orlando RC (2011) Role of E-cadherin in the pathogenesis of gastroesophageal reflux disease. Am J Gastroenterol 106: 1039–1047

21. Kadota T, Yano T, Kato T, Imajoh M, Noguchi M, Morimoto H, Osera S, Yoda Y, Oono Y, Ikematsu H, et al (2016) Prophylactic steroid administration for strictures after endoscopic resection of large superficial esophageal squamous cell carcinoma. Endosc Int Open 4: E1267–E1274

22. Kanai N, Yamato M, Ohki T, Yamamoto M & Okano T (2012) Fabricated autologous epidermal cell sheets for the prevention of esophageal stricture after circumferential ESD in a porcine model. Gastrointest Endosc 76: 873–881

23. Kandikattu HK, Venkateshaiah SU, Verma AK & Mishra A (2021) Tacrolimus (FK506) treatment protects allergen-, IL-5- and IL-13-induced mucosal eosinophilia. Immunology 163: 220–235

24. Kataoka M, Anzai S, Shirasaki T, Ikemiyagi H, Fujii T, Mabuchi K, Suzuki S, Yoshida M, Kawai T & Kitajima M (2015) Efficacy of short period, low dose oral prednisolone for the prevention of stricture after circumferential endoscopic submucosal dissection (ESD) for esophageal cancer. Endosc Int Open 3: E113–117

25. Kato R, Yamasaki Y & Tanaka S (2018) Triamcinolone injection and filling method to prevent stricture after esophageal endoscopic submucosal dissection. Dig Endosc 30: 795–796

26. Khlevner J, Jodorkovsky D, Bailey DD, Middlesworth W, Sethi A, Abrams J & Katzka DA (2023) Management of Adults With Esophageal Atresia. Clin Gastroenterol Hepatol 21: 15–25

27. Kiesslich R, Duckworth CA, Moussata D, Gloeckner A, Lim LG, Goetz M, Pritchard DM, Galle PR, Neurath MF & Watson AJM (2012) Local barrier dysfunction identified by confocal laser endomicroscopy predicts relapse in inflammatory bowel disease. Gut 61: 1146–1153

28. Kleuskens MTA, Haasnoot ML, Herpers BM, Ampting MTJ van, Bredenoord AJ, Garssen J, Redegeld FA & van Esch BCAM (2022) Butyrate and propionate restore interleukin 13-compromised esophageal epithelial barrier function. Allergy 77: 1510–1521

29. Koizumi E, Habu T, Goto O, Nakagome S, Ishikawa Y, Kirita K, Noda H, Higuchi K, Onda T, Akimoto T, et al (2023) Prophylactic use of steroids for a mucosal defect with a circumference of less than three-fourths prevents both symptomatic and asymptomatic stricture after esophageal endoscopic submucosal dissection. Surg Endosc 37: 5875–5882

30. Korbut E, Janmaat VT, Wierdak M, Hankus J, Wójcik D, Surmiak M, Magierowska K, Brzozowski T, Peppelenbosch MP & Magierowski M (2020) Molecular Profile of Barrett’s Esophagus and Gastroesophageal Reflux Disease in the Development of Translational Physiological and Pharmacological Studies. Int J Mol Sci 21: E6436

31. Kuo W-T, Zuo L, Odenwald MA, Madha S, Singh G, Gurniak CB, Abraham C & Turner JR (2021) The Tight Junction Protein ZO-1 Is Dispensable for Barrier Function but Critical for Effective Mucosal Repair. Gastroenterology 161: 1924–1939

32. Landin MD & Guerrón AD (2020) Endoscopic Mucosal Resection and Endoscopic Submucosal Dissection. Surg Clin North Am 100: 1069–1078

33. Letko E, Bhol K, Pinar V, Foster CS & Ahmed AR (1999) Tacrolimus (FK 506). Ann Allergy Asthma Immunol 83: 179–189; quiz 189–190

34. Li X, Mao M, Zhang Y, Yu K & Zhu W (2019) Succinate Modulates Intestinal Barrier Function and Inflammation Response in Pigs. Biomolecules 9: 486

35. Liu H, Zhang X, Liu Y, Xin N, Deng Y & Li Y (2022) Semen Ziziphi Spinosae attenuates blood– brain barrier dysfunction induced by lipopolysaccharide by targeting the FAK-DOCK180-Rac1- WAVE2-Arp3 signaling pathway. NPJ Sci Food 6: 27

36. Maxton DG, Ainley CC, Grainger SL, Morris RW & Thompson RP (1987) Teeth and benign oesophageal stricture. Gut 28: 61–63

37. Meyer W, Schoennagel B, Kacza J, Busche R, Hornickel IN, Hewicker-Trautwein M & Schnapper A (2014) Keratinization of the esophageal epithelium of domesticated mammals. Acta Histochem 116: 235–242

38. Moosbrugger-Martinz V, Jalili A, Schossig AS, Jahn-Bassler K, Zschocke J, Schmuth M, Stingl G, Eckl KM, Hennies HC & Gruber R (2015) Epidermal barrier abnormalities in exfoliative ichthyosis with a novel homozygous loss-of-function mutation in CSTA. Br J Dermatol 172: 1628–1632

39. Nagami Y, Ominami M, Shiba M, Sakai T, Fukunaga S, Sugimori S, Otani K, Hosomi S, Tanaka F, Taira K, et al (2018) Prediction of esophageal stricture in patients given locoregional triamcinolone injections immediately after endoscopic submucosal dissection. Dig Endosc 30: 198–205

40. Nakayama S, Sng N, Carretero J, Welner R, Hayashi Y, Yamamoto M, Tan AJ, Yamaguchi N, Yasuda H, Li D, et al (2014) β-catenin contributes to lung tumor development induced by EGFR mutations. Cancer Res 74: 5891–5902

41. Ohki T, Yamato M, Murakami D, Takagi R, Yang J, Namiki H, Okano T & Takasaki K (2006) Treatment of oesophageal ulcerations using endoscopic transplantation of tissue-engineered autologous oral mucosal epithelial cell sheets in a canine model. Gut 55: 1704–1710

42. Oyama T (2015) Prevention of stricture after large esophageal endoscopic submucosal dissections. Endoscopy 47: 289–290

43. Pernu H, Räsänen O, Salo T, Rinne A, Herva R & Järvinen M (1990) Cystatin A and B in the development of human squamous epithelia. Acta Histochem 88: 53–57

44. Puthia M, Storm P, Nadeem A, Hsiung S & Svanborg C (2014) Prevention and treatment of colon cancer by peroral administration of HAMLET (human α-lactalbumin made lethal to tumour cells). Gut 63: 131–142

45. Roh J-L, Lee J, Kim EH & Shin D (2017) Plasticity of oral mucosal cell sheets for accelerated and scarless skin wound healing. Oral Oncol 75: 81–88

46. van Roy F & Berx G (2008) The cell-cell adhesion molecule E-cadherin. Cell Mol Life Sci 65: 3756– 3788

47. Sakaguchi Y, Tsuji Y, Sato J, Kubota D, Obata M, Cho R, Nagao S, Miura Y, Ohki D, Mizutani H, et al (2023) Repeated steroid injection and polyglycolic acid shielding for prevention of refractory esophageal stricture. Surg Endosc 37: 6267–6277

48. Sakaguchi Y, Tsuji Y, Shinozaki T, Ohki D, Mizutani H, Minatsuki C, Niimi K, Yamamichi N & Koike K (2020) Steroid injection and polyglycolic acid shielding to prevent stricture after esophageal endoscopic submucosal dissection: a retrospective comparative analysis (with video). Gastrointest Endosc 92: 1176–1186.e1

49. Shi Q, Ju H, Yao L-Q, Zhou P-H, Xu M-D, Chen T, Zhou J-M, Chen T-Y & Zhong Y-S (2014) Risk factors for postoperative stricture after endoscopic submucosal dissection for superficial esophageal carcinoma. Endoscopy 46: 640–644

50. Shibagaki K, Ishimura N, Oshima N, Mishiro T, Fukuba N, Tamagawa Y, Yamashita N, Mikami H, Izumi D, Taniguchi H, et al (2018) Esophageal triamcinolone acetonide-filling method: a novel procedure to prevent stenosis after extensive esophageal endoscopic submucosal dissection (with videos). Gastrointest Endosc 87: 380–389

51. Takahashi H, Arimura Y, Okahara S, Kodaira J, Hokari K, Tsukagoshi H, Shinomura Y & Hosokawa M (2015) A randomized controlled trial of endoscopic steroid injection for prophylaxis of esophageal stenoses after extensive endoscopic submucosal dissection. BMC Gastroenterol 15: 1

52. Undre NA, Moloney FJ, Ahmadi S, Stevenson P & Murphy GM (2009) Skin and systemic pharmacokinetics of tacrolimus following topical application of tacrolimus ointment in adults with moderate to severe atopic dermatitis. Br J Dermatol 160: 665–669

53. Venkataramanan R, Swaminathan A, Prasad T, Jain A, Zuckerman S, Warty V, McMichael J, Lever J, Burckart G & Starzl T (1995) Clinical pharmacokinetics of tacrolimus. Clin Pharmacokinet 29: 404–430

54. Wallemacq PE & Reding R (1993) FK506 (tacrolimus), a novel immunosuppressant in organ transplantation: clinical, biomedical, and analytical aspects. Clin Chem 39: 2219–2228

55. Wang KK (2022) Endoscopic submucosal dissection and potential cancer dissemination. Gut 71: 236–237

56. Wang Q, Qi C, Min P, Wang Y, Ye F, Xia T, Zhang Y & Du J (2022) MICAL2 contributes to gastric cancer cell migration via Cdc42-dependent activation of E-cadherin/β-catenin signaling pathway. Cell Communication and Signaling : CCS 20

57. Wang Y, Xia W, Tian L, Zhu B, Chen M, Si X, Lin S & Gong Y (2023) Comparison of statins with steroids and botulinum toxin A in the prevention of benign strictures after esophageal endoscopic submucosal dissection: a retrospective cohort study. Surg Endosc

58. Wang Z, Zhou H, Zheng H, Zhou X, Shen G, Teng X, Liu X, Zhang J, Wei X, Hu Z, et al (2021) Autophagy-based unconventional secretion of HMGB1 by keratinocytes plays a pivotal role in psoriatic skin inflammation. Autophagy 17: 529–552

59. Wei R, Zhang L, Hu W, Wu J & Zhang W (2022) CSTA plays a role in osteoclast formation and bone resorption by mediating the DAP12/TREM2 pathway. Biochem Biophys Res Commun 627: 12–20

60. Xia S-Y, Lu Q, Wang Z-J, Gan T, Yang J-L & Wang Z (2023) Development and validation of a model to determine the risk of esophageal strictures after endoscopic submucosal dissection for esophageal neoplasms. Surg Endosc 37: 2163–2172

61. Yamaguchi N, Isomoto H, Nakayama T, Hayashi T, Nishiyama H, Ohnita K, Takeshima F, Shikuwa S, Kohno S & Nakao K (2011) Usefulness of oral prednisolone in the treatment of esophageal stricture after endoscopic submucosal dissection for superficial esophageal squamous cell carcinoma. Gastrointest Endosc 73: 1115–1121

62. Yeung BPM & Chiu PWY (2017) PTH-013 Retrospective study on oesophageal stricture after endoscopic submucosal dissection stricture in a regional tertiary referral centre. Gut 66: A211–A211

63. Zhang Q, Agoston AT, Pham TH, Zhang W, Zhang X, Huo X, Peng S, Bajpai M, Das K, Odze RD, et al (2019) Acidic Bile Salts Induce Epithelial to Mesenchymal Transition via VEGF Signaling in Non-Neoplastic Barrett’s Cells. Gastroenterology 156: 130–144.e10

64. Zhang X, Monnoye M, Mariadassou M, Beguet-Crespel F, Lapaque N, Heberden C & Douard V (2021) Glucose but Not Fructose Alters the Intestinal Paracellular Permeability in Association With Gut Inflammation and Dysbiosis in Mice. Front Immunol 12: 742584

65. Zhao Z, Zhao Q, Gu B, Yin C, Shen K, Tang H, Xia H, Zhang X, Zhao Y, Yang X, et al (2020) Minimally invasive implantation and decreased inflammation reduce osteoinduction of biomaterial. Theranostics 10: 3533–3545

